# Investigating the effects of prednisolone on behavior in mouse models of Duchenne muscular dystrophy

**DOI:** 10.1101/2024.10.29.620838

**Authors:** M.A.T. Verhaeg, D. van de Vijver, C.L. Tanganyika-de Winter, E.M. van der Pijl, L.J.M. Mastenbroek, U. Leka, T.L. Stan, M. van Putten

## Abstract

**Background:** Next to progressive muscle loss, Duchenne muscular dystrophy patients suffer from behavioral and cognitive problems. This is due to mutations in the *DMD* gene, that result in the lack of dystrophin in both the muscles and brain. As part of the standards of care, patients receive corticosteroids (prednisolone or deflazacort) to slow down muscle degeneration. The precise consequences of chronic corticosteroid usage on the behavior of DMD patients remain unclear, mainly due to challenges of recruiting corticosteroid naïve patients into clinical studies.

**Objective:** This study used DMD mouse models, representing mutations resulting in lack of one or more dystrophin isoforms, to analyze the effects of corticosteroid treatment on different behavioral domains.

**Methods:** Prednisolone (PDN) or placebo was administered via a subcutaneous 60-day slow release pellet (66 µg/day) and mice were subjected to several behavioral tests.

**Results:** Unfortunately, the pellet only exposed mice to PDN for half of the intended duration. During the time of PDN exposure, we found a small amelioration in anxiety but were unable to find any differences in social interaction and spatial learning and memory.

**Conclusions:** Short term exposure to PDN via a slow release pellet does not seem to negatively affect anxiety, social interaction or spatial learning and memory. We cannot rule out that a longer treatment period than 4 weeks would affect behavior in DMD mice.

## Introduction

Duchenne muscular dystrophy (DMD) is a progressive X-linked disorder that affects approximately 1 in 5,000 males (1). The disease is characterized by severe muscle wasting leading to loss of ambulation and eventually cardiac and respiratory failure resulting in premature death around the age of 30 to 40 years (2). Next to the muscle degeneration, approximately 30% of patients suffer from cognitive and behavioral problems, including autism spectrum disorder, obsessive-compulsive disorder, attention-deficit hyperactivity disorder, depression, anxiety, inattention, reading deficits and epilepsy (3-8). Parents report that these alterations in behavior and emotional functioning negatively impact the quality of life for both patients and their families (9, 10).

DMD is caused by mutations in the *DMD* gene, which normally encodes for the dystrophin protein. The *DMD* gene contains 7 promotor regions, resulting in multiple dystrophin isoforms which are expressed in different tissues and carry out diverse functions. One full-length dystrophin isoform (Dp427m) is expressed in muscle. The other full-length isoforms (Dp427c, Dp427p) and the shorter isoforms Dp140, Dp71 and Dp40 are expressed throughout the brain, but in particular, in the cortex, hippocampus and amygdala while lower levels are found in the cerebellum and pons (11-13). Depending on the location of the mutation in the *DMD* gene, patients either lack only the full-length dystrophin isoforms or one or multiple shorter isoforms in addition. Approximately 45% of patients lack only Dp427, 45% lack Dp427 and Dp140 and 3-10% lack all dystrophin isoforms (3, 14-17). The amount of missing isoforms seems to correlate with the incidence and severity of cognitive and behavioral problems (3, 14, 18).

The vast majority of DMD patients uses corticosteroids (prednisolone or deflazacort), either daily or in an intermittent regimen, as part of the standards of care to slow down the muscle degeneration (19-21). Although the chronic use positively affects muscle pathology, on the downside, it adversely affects patient’s weight and bone health and can cause short stature, behavioral changes, cataracts, hirsutism and cushingoid appearance (22). The changes in behavior are reported to be one of the most common reasons for the discontinuation of corticosteroid treatments among DMD patients (23, 24). In healthy individuals, acute corticosteroid use has led to increased negative emotions, poorer executive function and impaired short term and long term memory (25, 26). Chronic use of corticosteroids has negative effects on executive function, short term memory and concentration and can cause insomnia (26, 27). Research concerning the consequences of corticosteroid treatment on behavior and cognition in DMD patients has been very minimal. The low percentage of corticosteroid naïve DMD patients makes clinical studies challenging. Next to possible elevations in irritability (28), the type of behavioral problems was either not specified (29), or studies were unable to find any behavioral problems that are specifically caused, or affected in severity by the corticosteroid treatment in DMD patients (8, 30, 31). It has been reported that the type of treatment regimen can influence behavior. Prednisolone leads to more behavioral changes and more aggressive behavior when compared to deflazacort (32-35). Furthermore, a daily versus intermittent corticosteroid treatment regimen might also influence the effect on behavior, although reports are less consistent (3, 36). It is also known that the grey matter volume is more broadly and extensively altered in DMD patients receiving daily deflazacort treatment compared to intermittent prednisolone (37). Unfortunately, most studies did not elaborate on the type of behavioral changes that were observed, making it difficult to precisely understand which behavioral domains are being affected by corticosteroid treatment in DMD patients.

Mouse models could improve our understanding of the consequences of corticosteroid administration on behavior and cognition. In wildtype (WT) mice, corticosteroid treatment is associated with increased depression and anxiety (38-40), increased avoidance behavior (41), decreased food seeking behavior (42) and thereby reduced motivation in food rewarded tasks (43), and can interfere with learning and memory (44, 45). Knowledge on how corticosteroid treatment influences the brain of DMD mouse models remains limited. The most commonly used DMD model, the *mdx* mouse which lacks only the full length Dp427 isoforms, is described to exhibit increased anxiety and fear (46-51), altered social interaction (52) and deficits in long term memory (47, 53-56). *Mdx52* and *mdx*^*4cv*^ mice, lacking Dp140 in addition to Dp427, have similar types of deficits in terms of working memory and show a further decline in emotional reactivity, social interaction and fear learning compared to *mdx* mice (46, 57, 58). These behavioral alterations were all observed in corticosteroid naïve mice. *Mdx* mice treated for 28 days with a high dose of prednisolone show borderline increased depressive-like behavior, being detected in one out of two tests, but this altered behavior was not observed in mice treated with deflazacort (59). The open field test has been performed in corticosteroid treated and naïve mice. However, these studies mainly focused on locomotor activity, therefore the effects of corticosteroid treatment on anxious behavior remain unclear (60, 61). To this date, no studies on corticosteroid treatment have included mice lacking multiple dystrophin isoforms. Therefore, it remains unclear if the additional lack of Dp140 would lead to a different response to corticosteroid treatment compared to mice lacking only Dp427.

In this study, we aimed to unravel the effects of corticosteroid treatment on behavior in different DMD mouse models. Both *mdx* and *mdx*^*4cv*^ mice were included to represent the majority of isoform affecting mutations in DMD patients, which allowed us to compare possible differences in treatment response due to the additional lack of Dp140. Corticosteroid treatment was administered in the form of a subcutaneous prednisolone slow-release pellet. After implantation, *mdx* and *mdx*^*4cv*^ were exposed to a variety of behavioral assays to analyze anxiety, social interaction and learning and memory.

Unfortunately, due to technical issues with the pellets, rendering the pellets inactive after only half of the expected release time had elapsed, tests concerning recognition memory, spontaneous behavior, learning flexibility and fear response had to be excluded as they were performed outside of the pellet’s active release window. Mice treated with prednisolone showed somewhat less anxious behavior compared to placebo treated mice, although the effect was not consistent between tests. No differences were found in terms of social interaction or spatial learning, memory and reversal learning. Overall, the effects of prednisolone were very minimal in our mouse models in terms of behavioral changes.

## Materials and methods

### Mice

Male *mdx* (*mdx*(BL6); spontaneous mutation) (62), *mdx*^4cv^ (B6Ros.Cg-*Dmd*^*mdx-4Cv*^/J; generated with N-ethylnitrosourea (ENU) chemical mutagenesis) (63) and wildtype mice (C57BL/6J; WT), were bred at the animal facility of the Leiden University Medical Center (LUMC). Heterozygous females were paired with WT males to generate male DMD and WT littermates. WT mice for the experiments were taken equally from breeding couples of both DMD strains. Mice were housed in groups of 2 to 3 mice in individually ventilated cages (Makrolon type II) filled with sawdust and enriched with cardboard nesting (Bed-r’Nest BRN8SR) and bedding (LIGNOCEL BK-8-15-00433) material as well as an enrichment in the form of a cardboard tunnel (GLP fun tunnels mini 1022006). Mice had *ad libitum* access to water and standard RM3 chow (SDS, Essex, United Kingdom) and were kept with a 12 hour dark/light cycle. Experiments were performed between 07.00 and 18.00 at the animal facility of the LUMC in rooms dedicated for behavioral experiments. Timing of experiments was kept as similar as possible to minimize effects of circadian rhythm on behavior. Experiments were performed by 4 researchers, both male and female. The experiments were approved by the Animal Ethics Committee of the LUMC (AVD 1160020171407, PE.17.246.032, PE.17.246.053, PE.17306.02.013) and conform with the Directive 2010/63/EU of the European Parliament.

### Experimental setup

Four experimental groups were included in this study. In group A, the effect of PDN treatment on behavior was assessed. From the age of 7 weeks, *mdx* and *mdx*^*4cv*^ mice were treated with PDN or placebo, while WT mice were treated only with placebo pellets (n=16) (Fig. 1). Bodyweight was monitored twice a week. One week after the pellet implantation, behavioral experiments were started. At 59 days of age, mice were sacrificed by CO_2_ and levels of PDN markers were assessed by qPCR.

**Figure 1:**
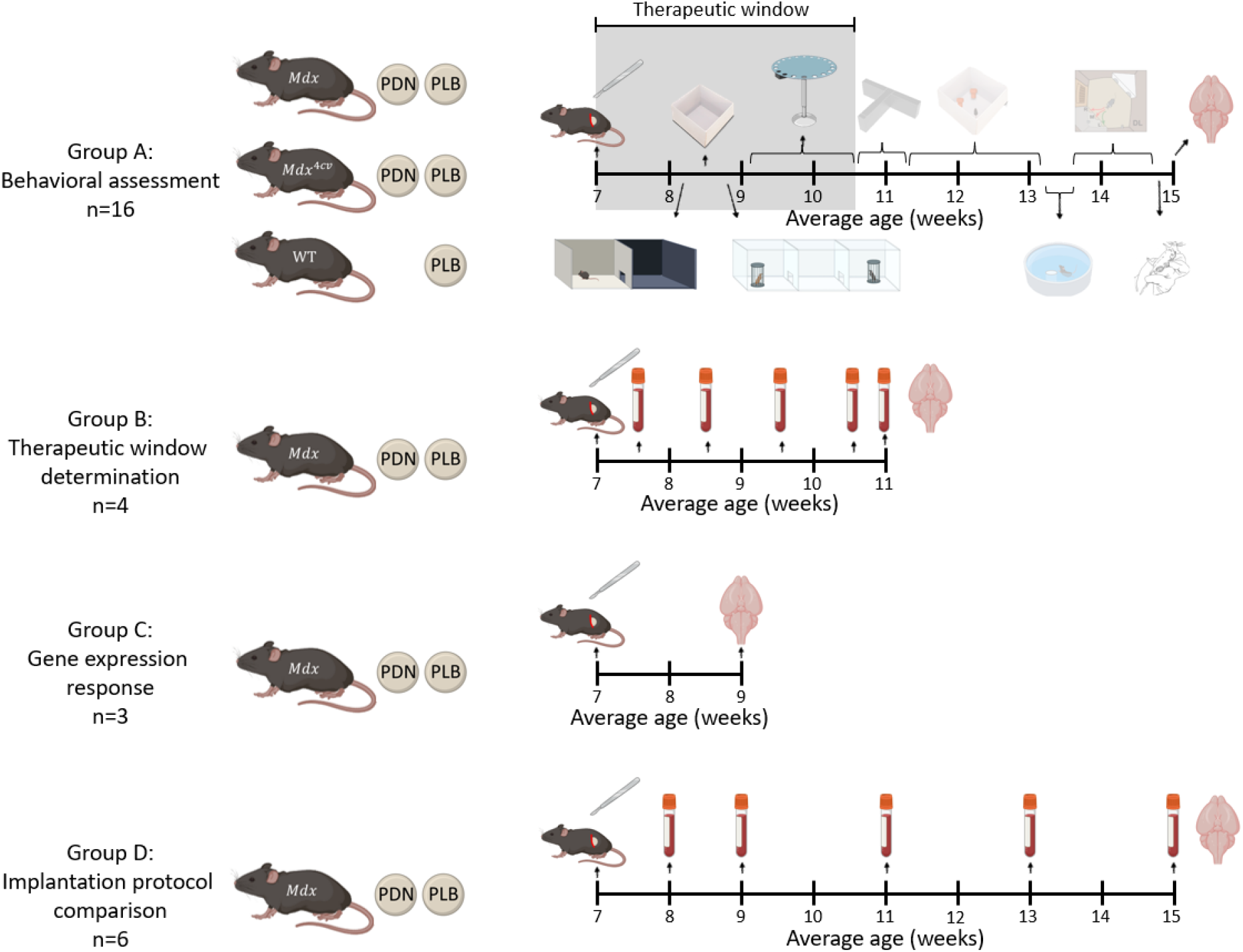
Study overview. Group A consisted of PDN or placebo (PLB) treated *mdx* and *mdx*^*4cv*^ mice, and placebo treated WTs (n=16/strain). After the pellet implantation at 7 weeks of age, animals could recover for 1 week before starting behavioral tests; the dark light box, the open field test, the 3 chamber social interaction test and the Barnes maze were performed during the therapeutic window. Other tests, including the T-maze, novel object recognition task, object placement task, Morris water maze, spontaneous behavior, discrimination and reversal learning and unconditioned fear test were performed outside the therapeutic window. At 15 weeks of age, 59 days after the pellet implantation, mice were sacrificed and the cortex, hippocampus and cerebellum were isolated for qPCR analysis of PDN marker genes. Group B consisted of PDN or placebo treated *mdx* mice only (n=4). After pellet implantation at 7 weeks of age, blood was drawn weekly until the point of sacrifice at 11 weeks of age for assessment of PDN levels. Cortex, hippocampus and cerebellum were isolated for qPCR analysis of PDN marker genes. Group C included *mdx* mice treated with PDN or placebo for 2 weeks (n=3), after which cortex, hippocampus and cerebellum were isolated for qPCR analysis of PDN marker genes. *Mdx* mice in group D received either a PDN pellet according to our implantation protocol, or a PDN or placebo pellet implanted according to the protocol of the pellet supplier, to assess if differences in implantation could influence the therapeutic window.

With group B, the active release window of the PDN pellets was determined. Hereto, *mdx* males were implanted with either a PDN or placebo pellet at 7 weeks of age (n=4). Blood was drawn weekly via an angled cut in the tail to assess PDN levels in the blood via an ELISA. After 4 weeks of treatment brains were collected and the levels of PDN markers were assessed.

With group C, the mode of action of PDN was tested. Male *mdx* mice were implanted with either a PDN or placebo pellet (n=3) and sacrificed 2 weeks after the start of treatment. Brains were collected to assess levels of PDN marker genes by qPCR.

With group D, two implantation protocols were compared to assess alterations in release time of the pellets. Male *mdx* mice were implanted with PDN pellets according to the protocol used during the earlier parts of the study, or with either a PDN or placebo pellets according to the protocol of the pellet supplier (n=6). Blood was drawn weekly via an angled cut in the tail to assess PDN blood levels via an ELISA and mice were sacrificed at 8 weeks after the pellet implantation.

### Pellet implantations

A prednisolone pellet (60 day slow-release, 4 mg PDN resulting in a daily dose of approximately 66 µg, Innovative Research of America, Sarasota, USA) or a placebo pellet from the same supplier was used to administer the treatment. Mice were injected with Temgesic (3 µl/g bodyweight, Indivior Europe Limited, Dublin 2, Ireland) 30 minutes before the pellet implantation to relieve pain. After induction with 4% isoflurane (0.5 L/min airflow), anesthesia was maintained at 2-2.5% isoflurane. Eye gel was applied (Addedpharma, Oss, The Netherlands) and mice were placed on a heating pad (37 °C). Skin on the left hip was shaved and cleaned with 70% ethanol before the pellet was implanted under the skin by making a small incision. The wound was closed using sutures (17 mm needle and Coated VICRYL thread (polyglactin 910) sutures, 95057-126, VWR).

The implantation protocol of the pellet supplier was comparable to the steps describe above, with the exception of the implantation site and the use of disinfectant; the pellet was implanted in the neck and no liquids, including ethanol, came into contact with the skin during the whole procedure.

### Blood collection

Mice were put under a heating lamp for 5 minutes prior to blood collection. Blood, 30-60 µl, was collected in a heparin coated tube (Sarstedt, Germany, Cat. No. 16.443) through an angled tail vein cut. Samples were spun down for 7 minutes at 13.000 rpm at 4 °C. Plasma was collected from the tubes and stored at -20 °C.

### Behavioral tests

Materials were cleaned with 70% ethanol between animals and trials. Animal behavior and location were tracked with Ethovision XT (Noldus, version 11.5), at a rate of 20 frames per second. Behavioral tests were performed between 8.00h and 16.00h, with a dedicated start time to keep variations in timing minimal between cohorts. All boxes used for behavioral testing were made in house. The order of the behavioral tests was chosen such to minimize the impact of the tests on each other.

#### Dark light box

To asses anxiety, mice were placed in a dark light box consisting of a dark and light compartment (each 50×25 cm) connected via a small latch (10×5 cm). The mouse was placed in the dark compartment for 20 seconds before the latch was opened. Thereafter, the mouse was able to freely move through both compartments for 5 minutes. Behavior was recorded in the light compartment only.

#### Open field

Mice were placed in the middle of the open field box (white, 50×50×35 cm) for 30 minutes. The box was virtually divided in an inner and outer zone, with the border at 10 cm from the walls.

#### 3 chamber social interaction test

To assess social preference, the 3 chamber social interaction test was performed. The opaque box (63×42×35 cm) was divided into 3 equal chambers by two transparent walls which had a small door (10×5 cm) to connect the chambers. The two lateral chambers contained a black tube (height; 20 cm high, diameter; 8 cm) with metal bars spaced 1 cm apart. The tube held an object, or a WT mouse that was unfamiliar to the experimental mouse (male, C57BL/6J, between 7 and 11 weeks old). A total of 5 WT pairs was used throughout the study. During habituation, both tubes were empty. Doors to the lateral chambers were closed and mice were placed in the middle compartment. After 2 minutes, the doors were opened simultaneously and the mice could explore for 10 minutes. Directly afterwards, the sociability trial was started. One of the tubes contained an object, while the other one contained an unfamiliar mouse. After the mouse was contained in the middle compartment for 30 seconds, doors were opened and the mice could explore freely for 10 minutes. During the next trial (social novelty seeking), the WT mouse that was used in the second trial was put into the same tube and compartment. The other tube contained an unfamiliar mouse. After being contained in the middle segment for 30 seconds, the experimental mice could explore freely again for 10 minutes. Interaction with the tubes was scored manually by two blinded scorers using Observer XT (Noldus, version 15).

#### Barnes maze

The Barnes maze consisted of a wooden circle (120 cm diameter) with 12 holes (10.5 cm diameter) equally spaced at 12 cm from the edge of the maze. A transparent platform underneath one of the holes led to a removable hidden escape box. Visual cues were hung around the maze for spatial orientation. During 5 acquisition days, mice were placed in the middle of the Barnes maze twice with a maximum interval of 5 minutes. Mice were allowed to explore the maze freely for a maximum of 5 minutes. If the mice did not find the escape box in time, they were put in the target hole. Mice were allowed to stay in the escape box for 30 seconds before being removed. During the second trial of the 5^th^ acquisition day, the platform and escape box were removed and the mice were placed in the middle of the maze and allowed to freely explore for 5 minutes. After two days of rest, mice underwent one acquisition day again consisting of two trials in which the platform and escape box were installed at the original position. The next day, the platform and escape box were moved to the opposite side of the maze. During this day, three trials were done with a maximum of 5 minutes in between them. The next day, two more trials were performed. During the first trial, the platform and escape box were still at this opposite side, while during the second trial both were removed. The interaction with the holes and the time spend till reaching the platform were scored manually by two blinded scorers using Observer XT. DeepLabCut was used to assess the distance walked until the moment of reaching the platform (64-66).

### Post-mortem analyses

#### RNA isolation cDNA synthesis and qPCR

Mice were sacrificed using CO_2_ (20-25% flow rate, 1 L/min CO_2_). Individual brain regions were isolated and snap frozen in liquid nitrogen and stored at -80 °C until further processing. RNA was isolated from the cortex, hippocampus and cerebellum (n=6 for group A, all brains for group B and C) using TRIsure (Bioline, London, United Kingdom). Tissue was disrupted using tubes filled with small beads (OPS Diagnostics, Lebanon, USA) in the MagNaLyzer and chloroform (Baker, FisherScience, Vantaa Finland) was added. Tubes were spun down, the upper phase was collected and 100% isopropanol (Baker) was added. After the tubes were spun down again, the supernatant was removed and the remaining pellet was resuspended in 100 µL RNase free H_2_O. RNA was further purified using RA1 buffer (Macherey-Nagel, Düren, Germany) and 100% ethanol. The lysate was loaded onto a NucleoSpin Column and spun down. Membrane desalting buffer was added and samples were spun down again. Reconstituted rDNase and reaction buffer (1:9) was added and the columns were incubated for 15 minutes at room temperature. The membrane was washed with RAW2 and RAW3 buffer and spun down. RNA was eluted in 40 µl RNase-free water. RNA concentrations were measured and 500 ng was taken and N6 primer was added before incubating for 10 minutes at 70 °C. A master mix of 5x first strand buffer, dNTPs and Tetro reverse transcriptase was made and added to the samples before incubation for 1h at 42 °C. cDNA was diluted 10x and stored at -20°C.

Expression levels of *Gilz, Mt2a* and *Fkbp5* were analyzed using qPCR to assess the mode of action of PDN in the brain. Two housekeeping genes were included; ribosomal protein L22 (*Rpl22*) and ribosomal protein L27 (*Rpl27*). Primer sequences can be found in Table S1. Per reaction, 4 µl 2x SensiMix Hi-ROX, 1 µl forward primer and 1 µl reverse primer (1 pmol/μL each) was added to 2 µl of sample. All samples were measured in triplo. Reverse transcriptase-negative and water controls were included. The 384-wells plate was centrifuged shortly at 2000 rpm and gene expression was assessed using the Light cycler (Roche) and analyzed with LinReg software. Gene expression was normalized against the housekeeping genes by dividing the individual sample values to the averaged concentration of the housekeeping genes.

#### Prednisolone ELISA

The ELISA was performed according to the manufacturer’s protocol of the ‘Mouse Prednisolone (PS) ELISA kit’ (MyBiosource, MBS265547, San diego, USA). In short, samples and a concentration curve were pipetted into the precoated plate and incubated at 37 °C for 90 minutes. Plates were washed repeatedly after every incubation. Antibodies were added and the plate was incubated for 60 minutes at 37 °C. After adding horseradish peroxidase (HRP)+ avidin, the plate was incubated at 37 °C for 30 minutes. Finally, substrates for the HRP reaction were added and after 30 minutes of incubation at 37°C, the last substrate was added to stop the reaction and sample concentrations were determined with a plate reader at 450 nm.

To compare the different implantation protocols (group D), serum samples of 2 mice within the same group were combined to reach sufficient volume for the ELISA (75 µl serum per duplo). For the other experimental groups samples were not combined.

### Data analysis

Statistical tests were performed using SPSS and R, while graphs were made with GraphPad Prism and Illustrator. Outcomes of WTs, taken from both *mdx* and *mdx*^*4cv*^ strains, were compared (Table S2) and since no differences were found, data was pooled before testing strain effects with *mdx* and *mdx*^*4cv*^ mutant mice. All data was assessed for normality and, if needed, transformed by log10 and additional statistical values were reported alongside the *P-*values (Table S3). Univariate two-way ANOVA tests for strain and treatment were performed with Tukey *post-hoc* tests for strain. To assess time differences in addition to strain and treatment, mixed linear models were used. All data is shown as average ± standard error of the mean (SEM). ^*^: *P*<0.05, ^**^: *P*<0.01, ^***^: *P*<0.001.

## Results

### Therapeutic window of the 60-day slow release PDN pellets was approximately 24 days

Due to the known side effect of PDN on the weight of mice, bodyweight was monitored twice a week. Weights of animals followed up to 59 days after implantation (group A) are depicted in Figure S1. After an initial drop of approximately 20% in bodyweight in PDN treated mice, values seemed to move back towards placebo levels at the end of the study. Surprisingly, even though the pellets were advertised as 60-day slow release pellets, expression of the PDN marker genes (*Gilz, Mt2a* and *Fkbp5*) was unchanged in the brains at 59 days after implantation (Fig. S2A). To evaluate the actual duration of the PDN exposure, we assessed PDN levels in serum with ELISA in a new cohort of *mdx* mice (group B). Blood PDN levels initially increased in response to the PDN treatment (*P* < 0.001) three days after the pellet implantation (Fig. 2A). However, between 17 and 28 days after implantation of the pellet, the PDN concentration dropped to placebo levels. In line, *Gilz* and *Mt2a* levels in the brain were also similar to placebo treated mice after 4 weeks of treatment (Figure S2B). To confirm the presence of PDN in the brain during the therapeutic window, expression of the PDN marker genes was analyzed in the brain of *mdx* mice 2 weeks after implantation (group C). Due to the small sample size, no statistics were performed, however, *Gilz* and *Mt2a* expression was higher in the PDN treated groups (Fig. 2B-D).

**Figure 2:**
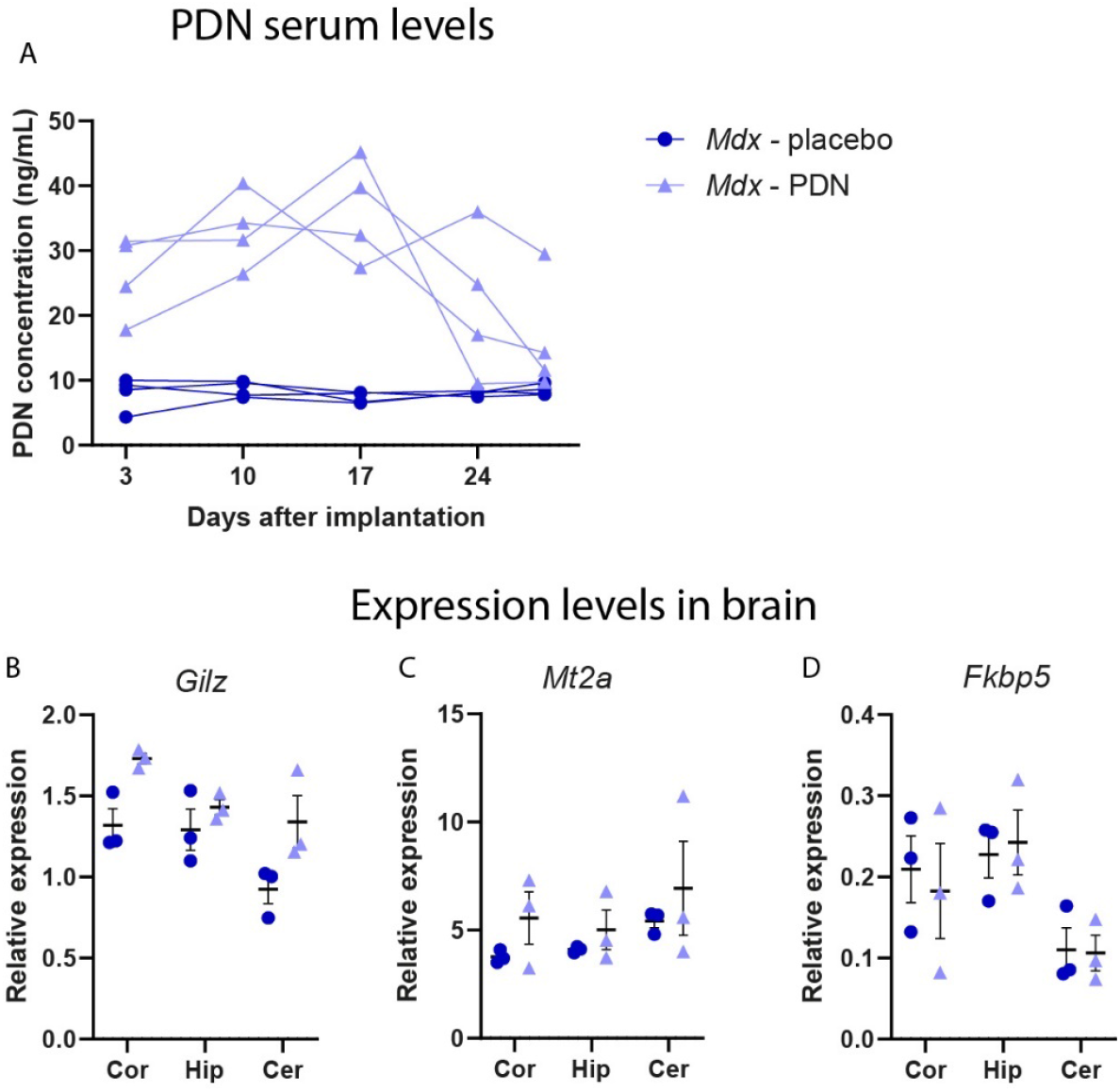
Therapeutic window of PDN pellets. A) ELISA analysis of PDN levels in serum samples of *mdx* mice. PDN treated mice showed a significant elevation in PDN levels until 24 days after implantation (*P* < 0.001). At 28 days after implantation, PDN levels were almost returned to placebo levels in 3 out of 4 animals. B-D) Expression levels of marker genes in response to PDN treatment in different brain regions assessed 2 weeks post pellet implantation. Elevations in *Gilz* and *Mt2a* expression can be seen, but not in *Fkbp5* expression. Note that no statistics were performed on the PDN marker gene expression levels due to the low sample size (n=3). Cor = cortex, Hip = hippocampus, Cer = cerebellum.

After consulting with the pellet supplier, they suggested several alterations to the implantation protocol which should lead to an active pellet release window of 60-days. We incorporated these changes in a new study directly comparing the effects of the implantation site (flank vs neck) and the influence of the use of skin disinfectant on the therapeutic window (group D). *Mdx* mice were implanted with a PDN pellet under both conditions and blood was collected after 1, 2, 4, 6 and 8 weeks to determine serum PDN levels. Notably, both implantation procedures led to a premature drop in PDN levels comparable to placebo within four weeks after implantation (Fig. S3). In conclusion, neither the implantation site nor the use or lack of disinfectant on the skin prior to implantation affected the performance of the pellets.

Even though behavioral tests were performed up to 7.5 weeks post pellet implantation in group A, based on these results, it was decided to only include behavioral tests performed during the therapeutic window (up to 24 days after the pellet implantation) in our analysis. These tests included the dark light box, the open field test, the 3 chamber social interaction task and the Barnes maze, respectively executed 8, 10, 12 and 15 to 24 days after implantation. Results of tests performed outside the therapeutic window (the T-maze, novel object recognition task, object placement task, Morris water maze, spontaneous behavior, discrimination and reversal learning and unconditioned fear) were excluded from the study.

### Slight decrease in anxious behavior due to PDN treatment

To assess the effects of genotype and PDN treatment on anxious behavior, mice were exposed to the dark light box and the open field test (Fig. 3). In the dark light box, *mdx* and *mdx*^*4cv*^ mice spent less time in total in the light compartment compared to WT mice (*P* = 0.006 & *P* < 0.001 respectively). This decreased tendency to explore the light compartment was stronger in *mdx*^*4cv*^ mice compared to *mdx* mice (*P* < 0.001) (Fig. 3A). The decreased motivation to explore the light compartment was also visible when calculating the average time spend in the compartment (Fig. 3B), as *mdx*^*4cv*^ spent less time in the compartment per visit compared to both WT and *mdx* mice (*P* = 0.009 & *P* = 0.003 respectively). Furthermore the average distance walked per visit was also decreased in *mdx*^*4cv*^ mice compared to both WT and *mdx* mice (both *P* < 0.001) (Fig. 3C). No effect of PDN treatment was observed in total time spend in the light compartment (Fig. 3A). Surprisingly, when the exploration time and distance walked were averaged per visit into the light compartment, PDN treated mice actually showed an increase in both time spend and distance walked in this department compared to placebo treated mice (*P* = 0.008 & *P* = 0.019 respectively) (Fig. 3B-C). However, no significant differences were found in the frequency of visits into the light compartment between PDN and placebo treated mice (data not shown).

**Figure 3:**
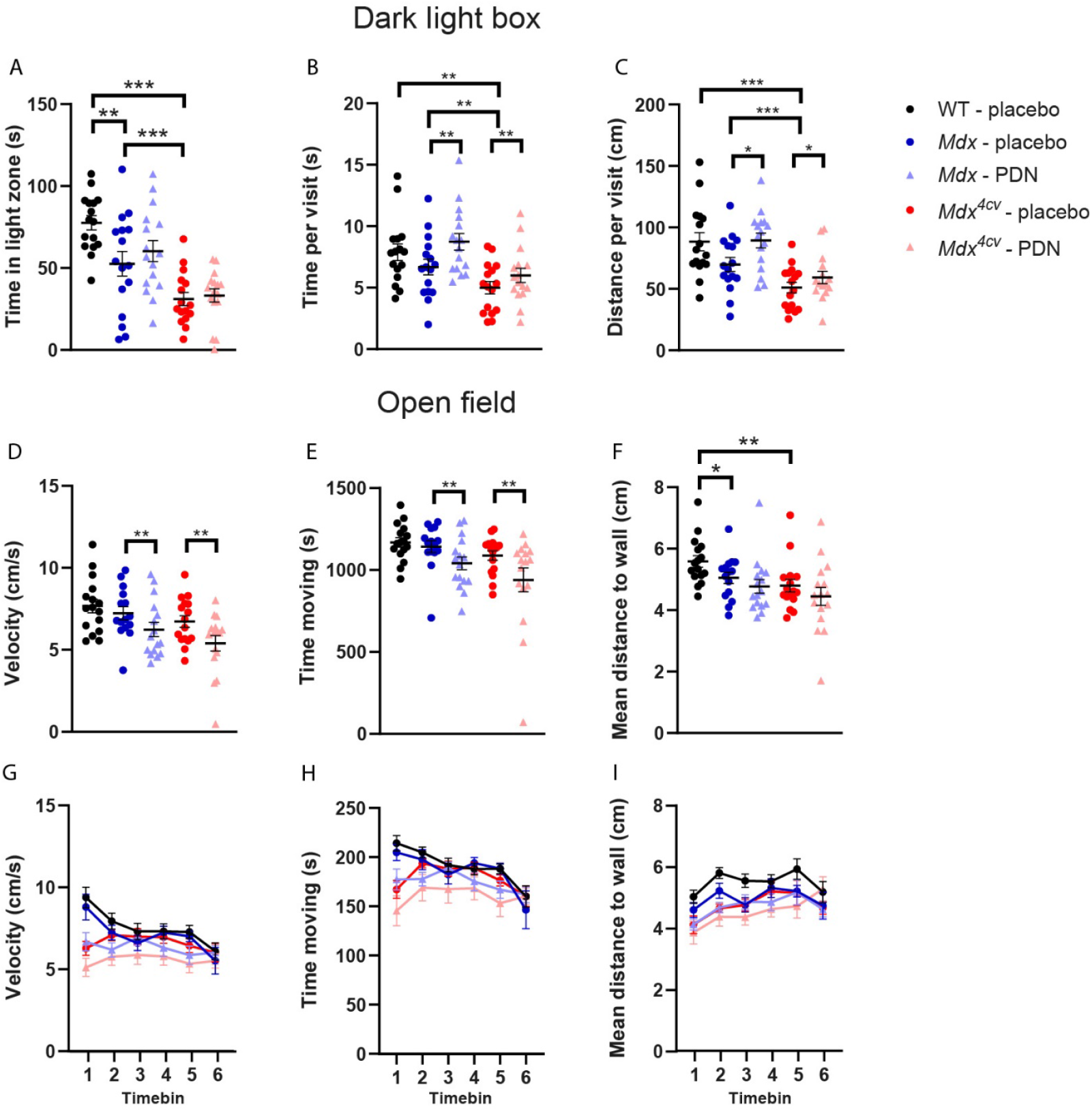
Anxious behavior in the dark light box and open field test. A) Total time in the light compartment was decreased in *mdx* mice compared to WTs (*P* = 0.006) and in *mdx*^*4cv*^ mice compared to both WT and *mdx* mice (both *P* < 0.001). B) When averaging the duration in the light compartment per visit, *mdx*^*4cv*^ mice spent less time in the light compartment compared to both WT and *mdx* mice (*P* 0.009 & *P* = 0.003 respectively). PDN treated mice spent on average more time in the light compartment compared to placebo treated mice (*P* = 0.008). C) *Mdx*^*4cv*^ mice walked less distance per visit in the light compartment compared to both WT and *mdx* mice (both *P* = 0.001). PDN treated mice walked more distance compared to placebo treated mice per visit into the light compartment (*P* = 0.019). D-E) In the open field test, PDN treated mice walked slower than placebo treated mice (*P* = 0.007) and spent less time moving (*P* = 0.006). F) Both *mdx* and *mdx*^*4cv*^ mice stayed closer to the walls during the open field test (*P* = 0.041 & *P* = 0.002 respectively). G-H) When analyzing the behavior during the open field task in time bins, *mdx*^*4cv*^ mice and PDN treated *mdx* mice showed an altered pattern in walking velocity and moving time during the first 5 minutes of the test. I) Average distance to the wall did not differ between groups when analyzed in time bins. ^*^: *P*<0.05, ^**^: *P*<0.01, ^***^: *P*<0.001.

During the open field test, no differences were found between strains in terms of locomotor activity, as seen by the velocity (Fig. 3D) and time spent moving (Fig. 3E). Both *mdx* and *mdx*^*4cv*^ mice stayed on average closer to the wall compared to WTs (*P* = 0.041 & *P* = 0.002 respectively) (Fig. 3F). PDN treated mice showed decreased locomotor function by walking slower and spending less time moving compared to placebo treated mice (*P* = 0.007 & *P* = 0.006) (Fig. 3D-E). However, anxious behavior, as indicated by the distance from the wall, was not altered in the PDN treated mice compared to placebo treated mice. Since the open field test was performed over a period of 30 minutes, parameters were also separated into five minute time bins to look for deviations in behavior over time (Fig. 3G-I). Interestingly, while both WT and placebo treated *mdx* mice showed a spike in velocity and time moved during the first 5 minutes of the open field test, this behavior was not seen in *mdx*^*4cv*^ mice (*P* = 0.043 & *P* = 0.019) or PDN treated *mdx* mice (*P* = 0.029 & *P* = 0.022). No differences in the distance to the wall were seen over time between any of the groups (Fig. 3I).

As expected, both DMD mouse models showed increased anxiety compared to WTs, with *mdx*^*4cv*^ mice showing slightly more anxious behavior compared to *mdx* mice. Overall, PDN treatment seemed to slightly decrease anxious behavior as indicated by the increased exploration of the light compartment in the dark light box, but this decrease in anxiety was not visible in the open field test. PDN treated mice showed decreased locomotor activity in the open field test, but no alterations in anxious behavior.

### PDN treatment does not affect sociability or social novelty seeking

To test social interaction in a relatively controlled setting, the 3 chamber social test was used (Fig. 4A). After habituation, mice could interact with either an object or a mouse in a tube (Fig. 4B). *Mdx*^*4cv*^ mice showed a preference towards interaction with the mouse in the tube compared to WTs (*P* = 0.025). No differences were found between PDN and placebo treated mice. Directly after the trial, the mice were reintroduced to the 3 chamber box, however, this time the object was switched with a novel mouse to test novel social interaction vs familiar social interaction (Fig. 4C). No differences were found in the discrimination index between strains or treatment groups. Taken together, while *mdx*^*4cv*^ mice show increased sociability, PDN treatment does not seem to affect social behavior in DMD mice.

**Figure 4:**
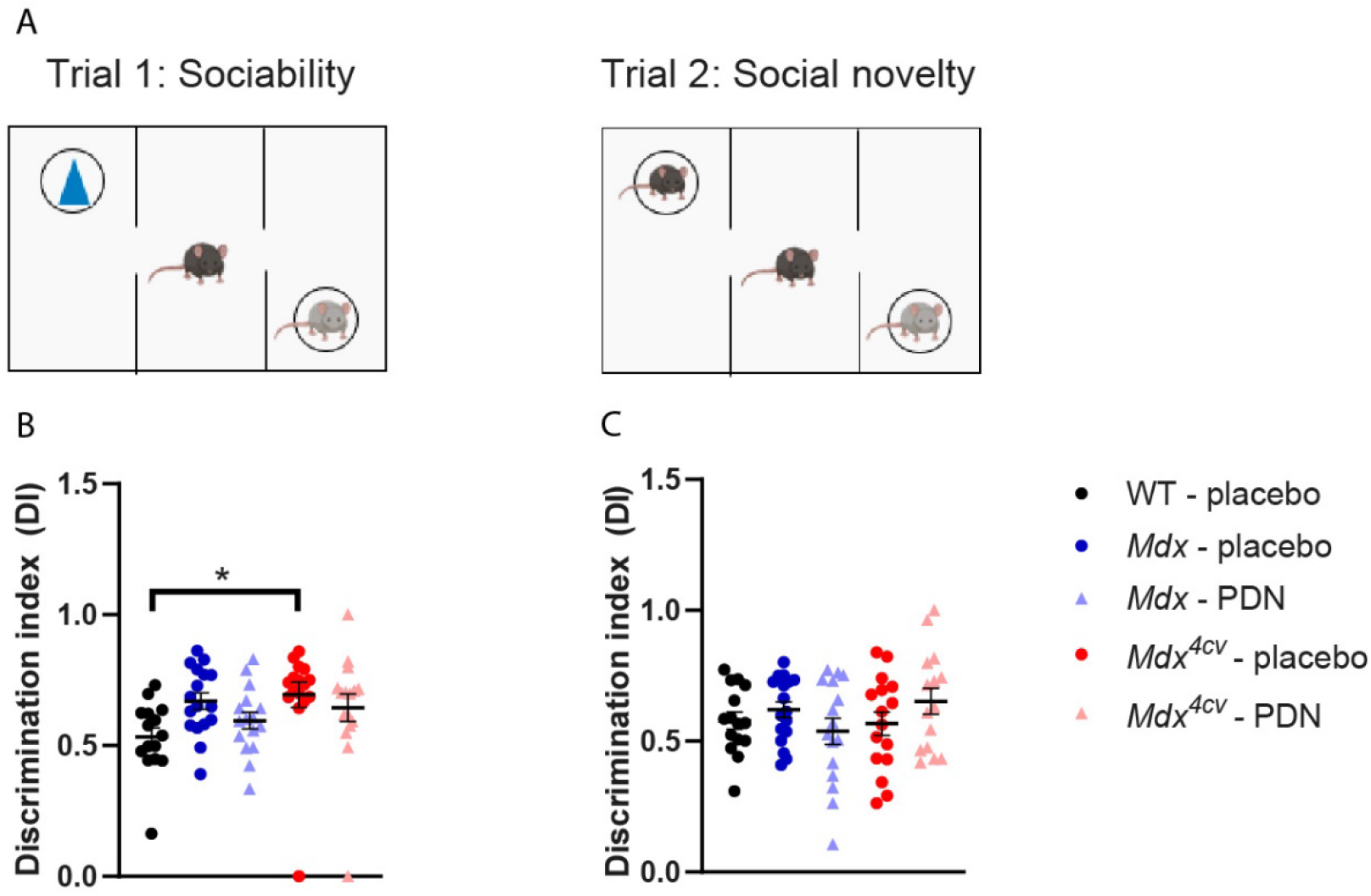
Sociability and social novelty seeking in the 3 chamber social interaction test. A) When given the choice between interaction with an object or an unfamiliar WT mouse, *mdx*^*4cv*^ mice showed a preference for social interaction compared to WTs (*P* = 0.025). B) No differences were found between groups in terms of preference for familiar versus novel social interaction. ^*^: *P*<0.05.

### Spatial learning, memory and reversal learning was not altered by PDN treatment

The Barnes maze was used to asses spatial learning, memory and reversal learning (Fig. 5A). Since velocity was decreased in *mdx*^*4cv*^ mice compared to WT and *mdx* mice (*P* = 0.049 & *P* = 0.031 respectively) (Fig. 5B), distance walked till reaching the target hole was used as a primary outcome measure instead of the more traditional latency to target hole. For five days, mice learned the location of the escape box. No differences were found in the distance mice needed to walk to reach the escape box during the learning (Fig. 5C), nor the probe test 24h later (Fig. 5D). Furthermore, no differences were found in the time the mice spent investigating the target hole during the probe test (Fig. 5E). During the reversal learning and the reversal probe, no differences were found in the distance walked to the target hole (Fig. 5F-G), nor in the interaction time with the target (Fig. 5H). Overall, spatial learning, memory and reversal learning does not seem to be altered by the lack of dystrophin nor by PDN treatment.

**Figure 5:**
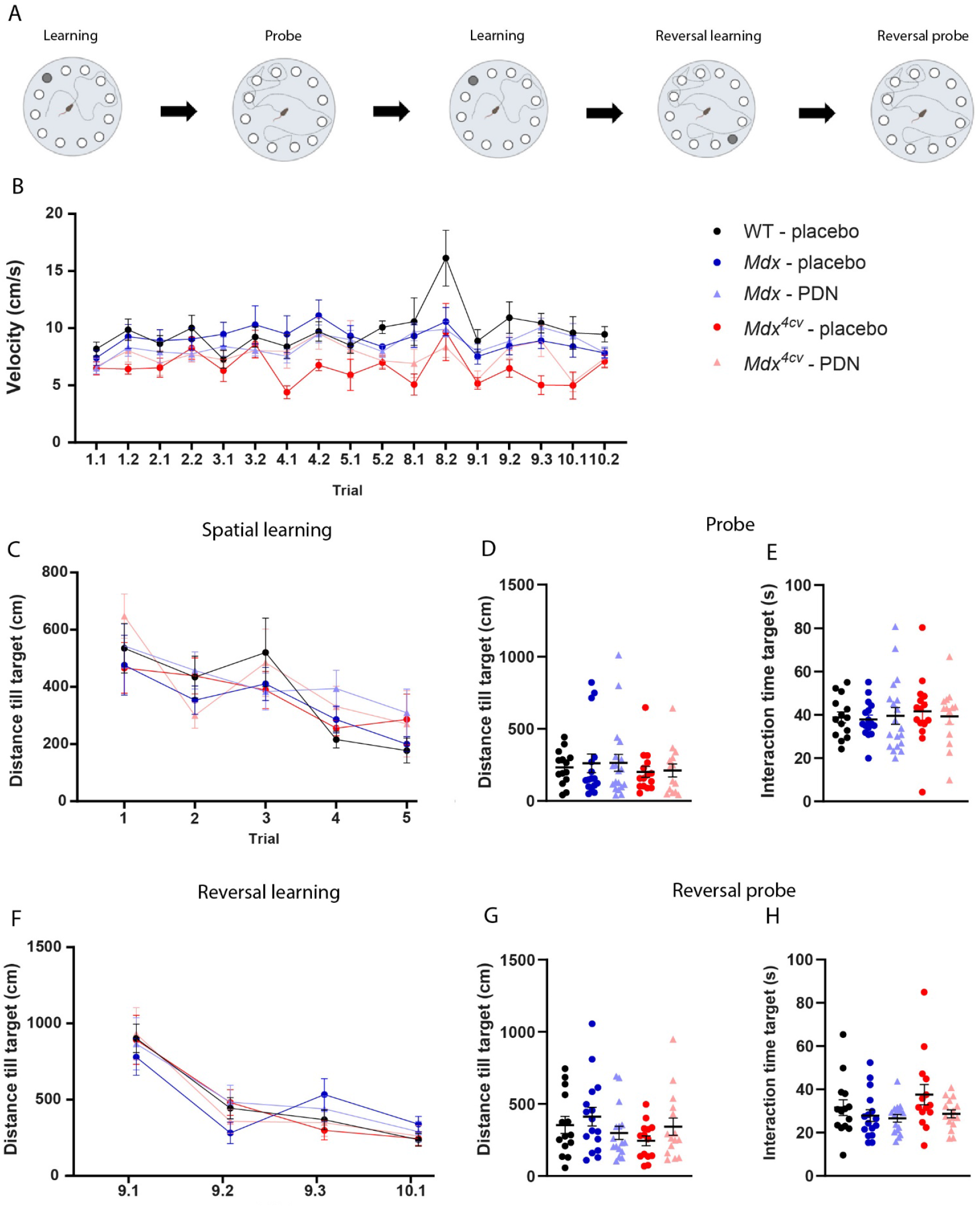
Spatial learning, memory and reversal learning in the Barnes maze. A) Schematic overview of the Barnes maze test and location of the escape box (in dark grey). B) Overall, *mdx* and *mdx*^*4cv*^ mice had a lower walking velocity compared to WT mice (*P* = 0.049 & *P* = 0.031 respectively). C-D) No differences were found between groups in terms of distance walked until reaching the target hole during the acquisition and the probe test. E) Interaction time with the target hole did not differ between groups. F-G) Distance walked till finding the new target location did not differ between groups during reversal learning or the reversal probe. H) Interaction time at the new target hole was similar between groups.

## Discussion

Corticosteroids are part of the standards of care in DMD and play an important role in slowing down the progression of the disease. Knowledge on the cognition and behavior related side effects are minimal, mainly due to the low number of corticosteroid naïve patients which hampers direct comparisons. Using two DMD mouse models, we aimed to assess the impact of chronic PDN usage on the brain and determine if the additional lack of Dp140 changes the influence of PDN on behavior.

Initially, multiple behavioral tests, assessing different domains, were conducted over a period of two months. Unfortunately, due to the shorter than expected release time of the PDN pellets, many results had to be excluded as they were obtained outside of the pellet’s therapeutic window. It should be noted that other research groups have had similar problems with the release time of other pellets from the same commercial supplier (67-71). None of those studies have investigated, reasoned or hypothesized as to why the pellets did not work for the expected duration. The supplier of the pellets stated that the use of ethanol to disinfect the skin before the implantation could have played a role in the shorter release time. Therefore, we repeated the experiment without exposing the skin to ethanol or any other liquid. However, we did not find any improvement in the duration of the therapeutic window. The reason behind this shortened therapeutic window remains unclear. At the endpoint, the pellets were still present in the mice, suggesting that they stopped dissolving prematurely, potentially due to encapsulation and that and none of the pellets had rapidly dissolved.

During the therapeutic window of the PDN pellets, only anxiety, social interaction and spatial learning, memory and reversal learning were assessed. Other tests, including a T-maze, Morris water maze, unconditioned fear test and object recognition and placement tasks were analyzed, but no differences were found between PDN and placebo treated mice. Since the tests fell outside the therapeutic window, results were not included in this study. Analyses of the tests performed in the PhenoTyper cages have therefore not been analyzed.

We found minimal or no effects of PDN on behavior. A slight decrease in anxiety was visible with the dark light box, in which during an average visit, the PDN treated mice showed increased exploration. This contradicts current literature, which reports increases in anxiety due to corticosteroid treatment (39, 40, 42, 72). However, since this effect was not visible in other parameters of the dark light box, nor in the open field test, this alteration in behavior does not seem very consistent or robust. PDN treated mice did show decreased activity in the open field test, which could be related to the weight loss of the animals. The open field test was performed 1.5 weeks after the pellet implantation, when the mice were experiencing a maximum weight loss of about 20%. This probably coincided with lower energy levels and therefore decreased overall activity as seen during the open field test. Notably, housing in ventilated cages can alter mouse behavior, especially in terms of depression (73). It is unclear if anxiety could also be affected by housing.

We were unable to find any effects of the PDN treatment on social interaction. Gasser *et al*. argues that social interaction, at least in the 3 chamber context, is controlled by androgen rather than by mineralocorticoids or glucocorticoids (74). Since our study used PDN, a glucocorticoid, this could explain our lack of differences found. It should be noted that there are many types of social interaction and that the context of the interaction and the characteristics of the social stimulus might influence the ability to detect differences. Saoudi *et al*. shows that this is the case for social deficits seen in *mdx52* mice (46). Possible deficits in social interaction, due to corticosteroid treatment could be overlooked due to the large variety of social interactions that have yet to be investigated. But for now, there are no indications that PDN influences social interaction in DMD mouse models.

No PDN-induced alterations were found in spatial learning, memory or reversal learning. However, since the Barnes maze was performed between day 14 and 24 after implantation, and PDN blood levels started to drop after 17 days, it could be possible that, especially during the reversal learning phase of the test, PDN levels in the brain were already too low to cause any alterations. Spatial learning and memory however, seem unaffected by the PDN treatment.

The overall lack of alterations due to PDN treatment was surprising as corticosteroids have been reported to affect anxiety and memory in WT mice (38-40, 44, 72). There are two possible explanations as to why this discrepancy with the literature could have occurred. Firstly, studies focusing on short term corticosteroid treatment (1 to 7 days of exposure), used very high concentrations of corticosteroids (up to 100 mg/kg) (42, 44), which are much more than what is used by DMD patients. The dose in our study, (66 µg/day) was lower than concentrations used in DMD patients. The dose was chosen as a maximal tolerable dose, as in a 7 day dose finding study, we observed that dosing of 33 µg/day already induced reductions in body weight (data not shown). Increasing the dose in future chronic studies could be harmful for the mice due to the excessive weight loss. The other difference between our study and most of the literature is the variation in treatment time. Most studies exposed mice to at least 3 weeks of corticosteroid treatment before assessing anxiety (38-40, 42, 72). While we also treated our mice for 3 weeks, the anxiety tests were performed 7 to 10 days after the treatment started. In terms of depression, Zhao *et al*. has shown that corticosteroids do not induce depressive like behavior after 6 days of treatment, but they did find a difference after 18 and 36 days of treatment, indicating the relevance of the duration of the corticosteroid treatment in depression (75). This could also apply when investigating the effects of corticosteroids on anxiety or learning and memory. Literature regarding learning and memory in mice is limited, but in rats, 90 days of PDN exposure of a clinically relevant dose induced deficits in spatial memory (76). DMD patients use corticosteroids for years, whereas our pellets only gave roughly 3 weeks of corticosteroid exposure and behavioral testing started already after 1 week. Probably, the exposure to PDN would have needed to be longer to allow detection of alterations in behavior. Therefore, it would be advisable to increase treatment duration before the start of behavioral assessments in the future.

Long term delivery of corticosteroids could be challenging in DMD mouse models. Daily injections or oral gavage are not favorable in DMD mouse models, since short periods of restraint induces strong freezing behavior which can last for hours even after multiple exposures. This fear reaction could influence the assessment of behavior (46, 48, 50, 51, 77, 78). Delivery via water or food seems a good alternative, as in WTs alterations on depression and anxiety could be identified regardless of the type of delivery (38, 40, 79-81). However, since prednisolone is unsolvable, another corticosteroid needs to be used in case of water delivery. Furthermore, the interval between treatments seems to influence the side effects on behavior in DMD patients (32-35). It remains unclear if this also applies to DMD mouse models. The type of delivery and the treatment interval should therefore be carefully considered when investigating the effects of corticosteroids in DMD.

Overall, PDN treatment via a slow release pellet did not seem to negatively impact anxiety, social interaction or spatial learning and memory after a short period of exposure. Studies further investigating the effects of PDN on the brain in DMD mouse models, utilizing different dosing regimens, are warranted in the future to gain a better understanding and eventually be able to counteract the negative side effects without losing the positive effects of the treatment.

## Acknowledgements

We would like to thank Esmée van der Linde and Kayleigh Putker for their help with sample processing, Dr. Luciano Censoni for help with the automated analysis of the Barnes maze data, Angel van Uffelen for her help with manual scoring of the 3 chamber test and Prof. Dr. Onno Meijer for his valuable advice.

## Data availability

The data supporting the findings of this study are available on request from the corresponding author.

## Ethical statement

The animal study was reviewed and approved by Central Authority for Scientific Procedures on Animals and performed according to Dutch regulation for animal experimentation, and in accordance with EU Directive 2010/63/EU.

## Author contributions

M.A.T. Verhaeg: Methodology, Investigation, Formal analysis, Writing – original draft, Writing – review and editing, Visualization

D. van de Vijver: Investigation

C.L. Tanganyika-de Winter: Investigation

E.M. van der Pijl: Investigation

L. Mastenbroek: Investigation

U. Leka: Investigation

T.L. Stan: Investigation

M. van Putten: Conceptualization, Methodology, writing-review and editing, supervision, funding acquisition

## Data availability

The data that support the findings of this study are available from the corresponding author upon request.

## References

1. Aartsma-Rus A, Ginjaar IB, Bushby K. The importance of genetic diagnosis for Duchenne muscular dystrophy. Journal of medical genetics. 2016.

2. Walter MC, Reilich P. Recent developments in Duchenne muscular dystrophy: facts and numbers. Journal of cachexia, sarcopenia and muscle. 2017;8(5):681.

3. Ricotti V, Mandy WP, Scoto M, Pane M, Deconinck N, Messina S, et al. Neurodevelopmental, emotional, and behavioural problems in Duchenne muscular dystrophy in relation to underlying dystrophin gene mutations. Developmental Medicine & Child Neurology. 2016;58(1):77–84.

4. Waite A, Brown SC, Blake DJ. The dystrophin–glycoprotein complex in brain development and disease. Trends in neurosciences. 2012;35(8):487–96.

5. Banihani R, Smile S, Yoon G, Dupuis A, Mosleh M, Snider A, et al. Cognitive and neurobehavioral profile in boys with Duchenne muscular dystrophy. Journal of child neurology. 2015;30(11):1472–82.

6. Hendriksen JG, Vles JS. Neuropsychiatric disorders in males with duchenne muscular dystrophy: frequency rate of attention-deficit hyperactivity disorder (ADHD), autism spectrum disorder, and obsessive—compulsive disorder. Journal of child neurology. 2008;23(5):477–81.

7. Hinton V, Fee R, Goldstein E, De Vivo D. Verbal and memory skills in males with Duchenne muscular dystrophy. Developmental Medicine & Child Neurology. 2007;49(2):123–8.

8. Hinton VJ, Nereo NE, Fee RJ, Cyrulnik SE. Social behavior problems in boys with Duchenne muscular dystrophy. Journal of Developmental & Behavioral Pediatrics. 2006;27(6):470–6.

9. Webb C. Parents’ perspectives on coping with Duchenne muscular dystrophy. Child: care, health and development. 2005;31(4):385–96.

10. Bendixen RM, Senesac C, Lott DJ, Vandenborne K. Participation and quality of life in children with Duchenne muscular dystrophy using the International Classification of Functioning, Disability, and Health. Health and quality of life outcomes. 2012;10:1–9.

11. Nudel U, Zuk D, Einat P, Zeelon E, Levy Z, Neuman S, et al. Duchenne muscular dystrophy gene product is not identical in muscle and brain. Nature. 1989;337(6202):76–8.

12. Holder E, Maeda M, Bies RD. Expression and regulation of the dystrophin Purkinje promoter in human skeletal muscle, heart, and brain. Human genetics. 1996;97:232–9.

13. Doorenweerd N, Mahfouz A, van Putten M, Kaliyaperumal R, t’Hoen PA, Hendriksen JG, et al. Timing and localization of human dystrophin isoform expression provide insights into the cognitive phenotype of Duchenne muscular dystrophy. Scientific reports. 2017;7(1):12575.

14. Taylor PJ, Betts GA, Maroulis S, Gilissen C, Pedersen RL, Mowat DR, et al. Dystrophin gene mutation location and the risk of cognitive impairment in Duchenne muscular dystrophy. PloS one. 2010;5(1):e8803.

15. Rasic MV, Vojinovic D, Pesovic J, Mijalkovic G, Lukic V, Mladenovic J, et al. Intellectual ability in the duchenne muscular dystrophy and dystrophin gene mutation location. Balkan Journal of Medical Genetics. 2014;17(2):25–35.

16. Desguerre I, Christov C, Mayer M, Zeller R, Becane H-M, Bastuji-Garin S, et al. Clinical heterogeneity of duchenne muscular dystrophy (DMD): definition of sub-phenotypes and predictive criteria by long-term follow-up. PloS one. 2009;4(2):e4347.

17. Pane M, Lombardo ME, Alfieri P, D’Amico A, Bianco F, Vasco G, et al. Attention deficit hyperactivity disorder and cognitive function in Duchenne muscular dystrophy: phenotype-genotype correlation. The Journal of pediatrics. 2012;161(4):705–9. e1.

18. Chamova T, Guergueltcheva V, Raycheva M, Todorov T, Genova J, Bichev S, et al. Association between loss of dp140 and cognitive impairment in duchenne and becker dystrophies. Balkan journal of medical genetics: BJMG. 2013;16(1):21.

19. Manzur AY, Kuntzer T, Pike M, Swan AV. Glucocorticoid corticosteroids for Duchenne muscular dystrophy. Cochrane database of systematic reviews. 2008(1).

20. Bushby K, Muntoni F, Urtizberea A, Hughes R, Griggs R. Treatment of Duchenne muscular dystrophy. Defining the gold standards of management in the use of corticosteroids. Report of the 124th ENMC International Workshop. Neuromuscul Disord. 2004;14:526–34.

21. Moxley III R, Ashwal S, Pandya S, Connolly A, Florence J, Mathews K, et al. Practice parameter: corticosteroid treatment of Duchenne dystrophy: report of the Quality Standards Subcommittee of the American Academy of Neurology and the Practice Committee of the Child Neurology Society. Neurology. 2005;64(1):13–20.

22. Gloss D, Moxley III RT, Ashwal S, Oskoui M. Practice guideline update summary: Corticosteroid treatment of Duchenne muscular dystrophy: Report of the Guideline Development Subcommittee of the American Academy of Neurology. Neurology. 2016;86(5):465–72.

23. Poysky J. Behavior patterns in Duchenne muscular dystrophy: report on the Parent Project Muscular Dystrophy behavior workshop 8–9 of December 2006, Philadelphia, USA. Neuromuscular Disorders. 2007;17(11-12):986–94.

24. Matthews DJ, James KA, Miller LA, Pandya S, Campbell KA, Ciafaloni E, et al. Use of corticosteroids in a population-based cohort of boys with duchenne and becker muscular dystrophy. Journal of child neurology. 2010;25(11):1319–24.

25. Schmidt LA, Fox NA, Goldberg MC, Smith CC, Schulkin J. Effects of acute prednisone administration on memory, attention and emotion in healthy human adults. Psychoneuroendocrinology. 1999;24(4):461–83.

26. Prado CE, Crowe SF. Corticosteroids and cognition: a meta-analysis. Neuropsychology Review. 2019;29(3):288–312.

27. Ciriaco M, Ventrice P, Russo G, Scicchitano M, Mazzitello G, Scicchitano F, et al. Corticosteroid-related central nervous system side effects. Journal of Pharmacology and Pharmacotherapeutics. 2013;4(1_suppl):S94–S8.

28. Angelini C. The role of corticosteroids in muscular dystrophy: a critical appraisal. Muscle & Nerve: Official Journal of the American Association of Electrodiagnostic Medicine. 2007;36(4):424–35.

29. Counterman KJ, Fatovic K, Good DC, Martin AS, Dasgupta S, Anziska Y. Associations Between Self-Reported Behavioral and Learning Concerns and DMD Isoforms in Duchenne Muscular Dystrophy. Journal of Neuromuscular Diseases. 2022;9(6):757–64.

30. Hendriksen JG, Poysky JT, Schrans DG, Schouten EG, Aldenkamp AP, Vles JS. Psychosocial adjustment in males with Duchenne muscular dystrophy: psychometric properties and clinical utility of a parent-report questionnaire. Journal of pediatric psychology. 2009;34(1):69–78.

31. Sienko S, Buckon C, Fowler E, Bagley A, Staudt L, Sison-Williamson M, et al. Prednisone and deflazacort in duchenne muscular dystrophy: do they play a different role in child behavior and perceived quality of life? PLoS currents. 2016;8.

32. Biggar WD, Skalsky A, McDonald CM. Comparing deflazacort and prednisone in Duchenne muscular dystrophy. Journal of neuromuscular diseases. 2022;9(4):463–76.

33. Griggs RC, Miller JP, Greenberg CR, Fehlings DL, Pestronk A, Mendell JR, et al. Efficacy and safety of deflazacort vs prednisone and placebo for Duchenne muscular dystrophy. Neurology. 2016;87(20):2123–31.

34. Balaban B, Matthews DJ, Clayton GH, Carry T. Corticosteroid treatment and functional improvement in Duchenne muscular dystrophy: long-term effect. American journal of physical medicine & rehabilitation. 2005;84(11):843–50.

35. Conway KC, Mathews KD, Paramsothy P, Oleszek J, Trout C, Zhang Y, et al. Neurobehavioral concerns among males with dystrophinopathy using population-based surveillance data from the muscular dystrophy surveillance, tracking, and research network. Journal of Developmental & Behavioral Pediatrics. 2015;36(6):455–63.

36. Escolar D, Hache L, Clemens P, Cnaan A, McDonald C, Viswanathan V, et al. Randomized, blinded trial of weekend vs daily prednisone in Duchenne muscular dystrophy. Neurology. 2011;77(5):444–52.

37. Geuens S, Van Dessel J, Govaarts R, Ikelaar NA, Meijer OC, Kan HE, et al. Comparison of two corticosteroid regimens on brain volumetrics in patients with Duchenne muscular dystrophy. Annals of Clinical and Translational Neurology. 2023;10(12):2324–33.

38. Gai BM, Bortolatto CF, Heck SO, Stein AL, Duarte MMMF, Zeni G, et al. An organoselenium compound improves behavioral, endocrinal and neurochemical changes induced by corticosterone in mice. Psychopharmacology. 2014;231:2119–30.

39. Lin L, Herselman MF, Zhou X-F, Bobrovskaya L. Effects of corticosterone on BDNF expression and mood behaviours in mice. Physiology & Behavior. 2022;247:113721.

40. Skupio U, Tertil M, Sikora M, Golda S, Wawrzczak-Bargiela A, Przewlocki R. Behavioral and molecular alterations in mice resulting from chronic treatment with dexamethasone: relevance to depression. Neuroscience. 2015;286:141–50.

41. Dieterich A, Srivastava P, Sharif A, Stech K, Floeder J, Yohn SE, et al. Chronic corticosterone administration induces negative valence and impairs positive valence behaviors in mice. Translational psychiatry. 2019;9(1):337.

42. Peng B, Xu Q, Liu J, Guo S, Borgland SL, Liu S. Corticosterone attenuates reward-seeking behavior and increases anxiety via D2 receptor signaling in ventral tegmental area dopamine neurons. Journal of neuroscience. 2021;41(7):1566–81.

43. Dieterich A, Stech K, Srivastava P, Lee J, Sharif A, Samuels BA. Chronic corticosterone shifts effort-related choice behavior in male mice. Psychopharmacology. 2020;237:2103–10.

44. Dos Santos N, Novaes LS, Dragunas G, Rodrigues JR, Brandão W, Camarini R, et al. High dose of dexamethasone protects against EAE-induced motor deficits but impairs learning/memory in C57BL/6 mice. Scientific reports. 2019;9(1):6673.

45. Schwabe L, Schächinger H, de Kloet ER, Oitzl MS. Corticosteroids operate as a switch between memory systems. Journal of cognitive neuroscience. 2010;22(7):1362–72.

46. Saoudi A, Zarrouki F, Sebrié C, Izabelle C, Goyenvalle Al, Vaillend C. Emotional behavior and brain anatomy of the mdx52 mouse model of Duchenne muscular dystrophy. Disease models & mechanisms. 2021;14(9):dmm049028.

47. Remmelink, Aartsma-Rus A, Smit A, Verhage M, Loos M, van Putten M. Cognitive flexibility deficits in a mouse model for the absence of full-length dystrophin. Genes, Brain and Behavior. 2016;15(6):558–67.

48. Vaillend C, Chaussenot R. Relationships linking emotional, motor, cognitive and GABAergic dysfunctions in dystrophin-deficient mdx mice. Human molecular genetics. 2017;26(6):1041–55.

49. Manning J, Kulbida R, Rai P, Jensen L, Bouma J, Singh SP, et al. Amitriptyline is efficacious in ameliorating muscle inflammation and depressive symptoms in the mdx mouse model of Duchenne muscular dystrophy. Experimental physiology. 2014;99(10):1370–86.

50. Razzoli M, Lindsay A, Law ML, Chamberlain CM, Southern WM, Berg M, et al. Social stress is lethal in the mdx model of Duchenne muscular dystrophy. EBioMedicine. 2020;55.

51. Sekiguchi M, Zushida K, Yoshida M, Maekawa M, Kamichi S, Yoshida M, et al. A deficit of brain dystrophin impairs specific amygdala GABAergic transmission and enhances defensive behaviour in mice. Brain. 2009;132(1):124–35.

52. Miranda R, Nagapin F, Bozon B, Laroche S, Aubin T, Vaillend C. Altered social behavior and ultrasonic communication in the dystrophin-deficient mdx mouse model of Duchenne muscular dystrophy. Molecular autism. 2015;6:1–17.

53. Bagdatlioglu E, Porcari P, Greally E, Blamire AM, Straub VW. Cognitive impairment appears progressive in the mdx mouse. Neuromuscular Disorders. 2020;30(5):368–88.

54. Vaillend C, Billard J-M, Laroche S. Impaired long-term spatial and recognition memory and enhanced CA1 hippocampal LTP in the dystrophin-deficient Dmdmdx mouse. Neurobiology of disease. 2004;17(1):10–20.

55. Sesay A, Errington M, Levita L, Bliss T. Spatial learning and hippocampal long-term potentiation are not impaired in mdx mice. Neuroscience letters. 1996;211(3):207–10.

56. Comim CM, Ventura L, Freiberger V, Dias P, Bragagnolo D, Dutra ML, et al. Neurocognitive impairment in mdx mice. Molecular neurobiology. 2019;56:7608–16.

57. Hashimoto Y, Kuniishi H, Sakai K, Fukushima Y, Du X, Yamashiro K, et al. Brain Dp140 alters glutamatergic transmission and social behaviour in the mdx52 mouse model of Duchenne muscular dystrophy. Progress in Neurobiology. 2022;216:102288.

58. Verhaeg M, Adamzek K, van de Vijver D, Putker K, Engelbeen S, Wijnbergen D, et al. Learning, memory and blood–brain barrier pathology in Duchenne muscular dystrophy mice lacking Dp427, or Dp427 and Dp140. Genes, Brain and Behavior. 2024;23(3):e12895.

59. Liu G, Lipari P, Mollin A, Jung S, Teplova I, Li W, et al. Comparison of pharmaceutical properties and biological activities of prednisolone, deflazacort, and vamorolone in DMD disease models. Human molecular genetics. 2024;33(3):211–23.

60. Guerron AD, Rawat R, Sali A, Spurney CF, Pistilli E, Cha H-J, et al. Functional and molecular effects of arginine butyrate and prednisone on muscle and heart in the mdx mouse model of Duchenne Muscular Dystrophy. PloS one. 2010;5(6):e11220.

61. Sali A, Guerron AD, Gordish-Dressman H, Spurney CF, Iantorno M, Hoffman EP, et al. Glucocorticoid-treated mice are an inappropriate positive control for long-term preclinical studies in the mdx mouse. PloS one. 2012;7(4):e34204.

62. Veltrop M, van der Kaa J, Claassens J, van Vliet L, Verbeek S, Aartsma-Rus A. Generation of embryonic stem cells and mice for duchenne research. PLoS currents. 2013;5.

63. Chapman VM, Miller DR, Armstrong D, Caskey CT. Recovery of induced mutations for X chromosome-linked muscular dystrophy in mice. Proceedings of the National Academy of Sciences. 1989;86(4):1292–6.

64. Forys B, Xiao D, Gupta P, Boyd JD, Murphy TH. Real-time markerless video tracking of body parts in mice using deep neural networks. bioRxiv. 2018:482349.

65. Mathis A, Mamidanna P, Cury KM, Abe T, Murthy VN, Mathis MW, et al. DeepLabCut: markerless pose estimation of user-defined body parts with deep learning. Nature neuroscience. 2018;21(9):1281–9.

66. Nath T, Mathis A, Chen AC, Patel A, Bethge M, Mathis MW. Using DeepLabCut for 3D markerless pose estimation across species and behaviors. Nature protocols. 2019;14(7):2152–76.

67. Hashim PH, Kinnear HM, Cruz CD, Padmanabhan V, Moravek MB, Shikanov A. Pharmacokinetic comparison of three delivery systems for subcutaneous testosterone administration in female mice. General and comparative endocrinology. 2022;327:114090.

68. Strom JO, Theodorsson E, Holm L, Theodorsson A. Different methods for administering 17β-estradiol to ovariectomized rats result in opposite effects on ischemic brain damage. BMC neuroscience. 2010;11:1–9.

69. Ingberg E, Theodorsson A, Theodorsson E, Strom J. Methods for long-term 17β-estradiol administration to mice. General and comparative endocrinology. 2012;175(1):188–93.

70. Robinson SA, Brookshire BR, Lucki I. Corticosterone exposure augments sensitivity to the behavioral and neuroplastic effects of fluoxetine in C57BL/6 mice. Neurobiology of stress. 2016;3:34–42.

71. Kott J, Mooney-Leber S, Shoubah F, Brummelte S. Effectiveness of different corticosterone administration methods to elevate corticosterone serum levels, induce depressive-like behavior, and affect neurogenesis levels in female rats. Neuroscience. 2016;312:201–14.

72. Odland AU, Sandahl R, Andreasen JT. Chronic corticosterone improves perseverative behavior in mice during sequential reversal learning. Behavioural Brain Research. 2023;450:114479.

73. York JM, McDaniel AW, Blevins NA, Guillet RR, Allison SO, Cengel KA, et al. Individually ventilated cages cause chronic low-grade hypoxia impacting mice hematologically and behaviorally. Brain, behavior, and immunity. 2012;26(6):951–8.

74. Gasser BA, Kurz J, Senn W, Escher G, Mohaupt MG. Stress-induced alterations of social behavior are reversible by antagonism of steroid hormones in C57/BL6 mice. Naunyn-Schmiedeberg’s archives of pharmacology. 2021;394:127–35.

75. Zhao Y, Xie W, Dai J, Wang Z, Huang Y. The varying effects of short-term and long-term corticosterone injections on depression-like behavior in mice. Brain research. 2009;1261:82–90.

76. Ramos-Remus C, González-Castañeda RE, González-Perez O, Luquin S, García-Estrada J. Prednisone induces cognitive dysfunction, neuronal degeneration, and reactive gliosis in rats. Journal of investigative medicine. 2002;50(6):458–64.

77. Yamamoto K, Yamada D, Kabuta T, Takahashi A, Wada K, Sekiguchi M. Reduction of abnormal behavioral response to brief restraint by information from other mice in dystrophin-deficient mdx mice. Neuromuscular Disorders. 2010;20(8):505–11.

78. Lindsay A, Russell AP. The unconditioned fear response in dystrophin-deficient mice is associated with adrenal and vascular function. Scientific Reports. 2023;13(1):5513.

79. Camargo A, Dalmagro AP, Platt N, Rosado AF, Neis VB, Zeni ALB, et al. Cholecalciferol abolishes depressive-like behavior and hippocampal glucocorticoid receptor impairment induced by chronic corticosterone administration in mice. Pharmacology Biochemistry and Behavior. 2020;196:172971.

80. Demuyser T, Deneyer L, Bentea E, Albertini G, Van Liefferinge J, Merckx E, et al. In-depth behavioral characterization of the corticosterone mouse model and the critical involvement of housing conditions. Physiology & Behavior. 2016;156:199–207.

81. van Donkelaar EL, Vaessen KR, Pawluski JL, Sierksma AS, Blokland A, Cañete R, et al. Long-term corticosterone exposure decreases insulin sensitivity and induces depressive-like behaviour in the C57BL/6NCrl mouse. PLoS One. 2014;9(10):e106960.

